# Cognitive Performance and Brain-Predicted Age Difference in Bipolar Disorder

**DOI:** 10.1101/2025.11.20.681142

**Authors:** Hui Xin Ng, Ashley Sutherland, Lisa Eyler

## Abstract

Neuroimaging-derived brain-predicted age difference (brain-PAD) is a promising marker of advanced brain aging, but its link to cognitive function in bipolar disorder (BD) is not well understood, especially when comparing across publicly available algorithms trained on diverse, large sample datasets and to algorithms trained on local cohorts with rich multimodal imaging data. Our study compares algorithms used to estimate brain-PAD in terms of their clinical relevance to cognition in BD. We included 44 euthymic BD I individuals and 73 HCs who completed the Delis-Kaplan Executive Function System, and we selected nine scores from this battery for further analyses. Raw scores were log-transformed, scaled, and subjected to PCA; PC1 indexed overall executive function. Four brain-PAD algorithms (PHOTON, BrainageR, DenseNet, Multimodal) were applied to T1-weighted MRI data; the multimodal algorithm also included Diffussion Tensor Imaging (DTI), Arterial Spin Labeling (ASL), functional Magnetic resonance imaging (fMRI) and resting state Magnetic resonance imaging (rsMRI) data. For each algorithm, we regressed brain-PAD on age, sex, and their interaction to obtain residuals, then used those residualized brain-PADs (which we refer to subsequently as brain-PADs throughout the text) to predict PC1. We then directly assessed if there were group differences in the relationship of brain-PAD to cognitive function by including an interaction term between group x brain-PAD. We found no significant group x brain-PAD interaction across all four algorithms. Given that, we then combined BD and HC and explored whether brain-PAD was a meaningful predictor of cognitive performance. Multimodal brain-PAD emerged as a strong negative predictor of cognitive performance (Beta Estimate = -0.084, SE = 0.024, t = -3.50, p < 0.001), indicating that those with older-appearing brains, as indexed by the brain-PAD, scored lower on PC1. BrainageR brain-PAD also significantly predicted PC1 (Beta Estimate = -0.031, SE = 0.0116, t = -2.71, p < 0.01), and DenseNet brain-PAD showed a modest effect (Beta Estimate = -0.0355, SE = 0.0177, t = -2.00, p < 0.05). PHOTON brain-PAD demonstrated a negative trend with PC1 (Beta Estimate = -0.024, SE = 0.0127, t = -1.92, p = 0.06). Residualized brain-PAD, after accounting for age and sex, was inversely associated with a composite metric of executive functioning, particularly for an algorithm integrating a range of imaging modalities. Our findings demonstrate how brain aging patterns captured by a neuroimaging-based, ML-derived composite metric could be associated with cognitive performance across algorithms trained on a variety of data granularity and sample sizes.

## Introduction

Brain-predicted age difference (brain-PAD) scores are calculated by subtracting chronological age from ‘brain’ age, which is estimated using neuroimaging data. Higher scores reflect advanced aging and are associated with increased mortality risk and poorer physical functioning (Cole 2018). Prior work has demonstrated higher brain-PAD in adults was negatively correlated with performance on measures of general cognitive status, processing speed, visual attention, and cognitive flexibility (Boyle et al 2021, Baecker et al 2021). One study that estimated brain-PAD in a sample of 869 45-year-old adults found that the individuals with higher brain-PAD tended to have poorer cognitive function in both adulthood and childhood, as well as impaired brain health as early as age three; findings from this study suggests that brain-PAD may also capture neurodevelopmental dysfunction present since childhood (Elliott et al 2021). Individuals with higher brain-PAD exhibit lower full-scale IQ and subscale scores in domains such as verbal comprehension, processing speed, working memory, and executive function (Elliott et al 2021).

Prior studies applying brain age algorithms in BD have reported mixed results— ranging from no group differences (Hajek et al., 2019; Nenadić et al., 2017; Shahab et al., 2019), to modest brain-PAD increases (Tønnesen et al., 2020), or lithium-related reductions (Van Gestel et al., 2019). A meta-analysis reported a small overall increase in BD (Ballester et al., 2022). As noted in Chapter 1, the ENIGMA-BD mega-analysis (Ng et al., n.d.) found brain-PAD increased with age in BD, but not with diagnosis alone. However, no prior BD-focused studies have examined links between brain-PAD and cognition.have higher brain-PAD as age increases, although there was no main effect of BD status.

Demro et al (2022) found that individuals with psychotic features and diagnosed with schizophrenia (SZ) or BD I exhibit an advanced brain-PAD of 5-8 years relative to HCs; larger brain-PADs were also linked to lower IQ and global assessment scores, especially among first-degree relatives. Lower cognitive performance predicts increased brain-PAD, which supports a neurodevelopmental rather than neurodegenerative origin of structural brain abnormalities (Demro et al., 2022). Salisbury et al (2024) found that lower predicted brain age was associated with higher vocabulary performance, which has been shown to be relatively robust against the effects of psychosis on cognition; however, they found no differences in the predicted brain age between first-episode SZ spectrum individuals and HCs.

Bipolar disorder (BD) is characterized by notable cognitive dysfunction; executive functioning has also been consistently identified as a key area of impairment. Deficits in working memory often persist even during euthymic periods and can significantly impact daily functioning (Jones et al.,2022) and older adults with BD showed a steeper decline in executive function than older HCs (Seelye et al., 2019). We chose to focus on executive functioning as prior studies have supported the relevance of the nine selected D-KEFS scores in capturing meaningful dimensions of executive functioning. For example, Savla et al. (2012) found that Trail Making and Stroop-based tasks clustered into distinct factors reflecting cognitive flexibility and abstraction, even after accounting for general processing speed. Similarly, Clark et al. (2010) identified separable components of inhibition/set-shifting and mental flexibility using PCA on D-KEFS subtests in individuals with schizophrenia.

Konstantinopoulou et al. (2021) demonstrated that Stroop and Trail Making performance loaded on distinct executive function domains—namely inhibition and set-shifting—in a mixed clinical sample. Finally, Camilleri et al. (2021) applied machine learning–based matrix factorization and PCA to D-KEFS data in healthy adults and identified a robust two-factor structure distinguishing lower-level speeded sequencing tasks (like TMT-A and Stroop Color Naming) from more complex executive processes (like inhibition and switching). Together, these findings support the relevance and interpretability of the selected D-KEFS scores in capturing meaningful and dissociable dimensions of executive function.

Examining the relationship between brain-PAD and executive function in BD and HC is important for validating brain-PAD as a meaningful biomarker for cognitive decline in BD. Chakrabarty et al. (2022) observed higher brain-PAD in first-episode bipolar disorder, with lower brain-PAD linked to poorer cognitive performance in an adolescent cohort, suggesting a delay relative to normative neurodevelopmental processes. However, it remains unclear which brain age algorithms have the highest sensitivity to BD-related alterations in cognition and the brain. Shared sensitivity to cognitive decline across algorithms would support brain-PAD as a robust biomarker for cognitive decline in BD, while discrepancies between algorithms could indicate that some are more sensitive to BD-related cognitive dysfunction than others. Our study helps identify the most reliable brain age algorithms for tracking cognitive decline in BD. Understanding which algorithms are more sensitive to BD-related cognitive dysfunction can improve the precision of brain-PAD as a biomarker, guiding more targeted interventions and better clinical decision-making for individuals with BD.

Our study compares four brain age prediction algorithms: PHOTON (Ridge Regression; L. K. M. Han, Dinga, et al., 2021), brainageR (Gaussian Process Regression), DenseNet (Wood et al., 2022), and Multimodal (Ridge Regression) to assess the sensitivity of these algorithms group differences in executive functioning. More specifically, we evaluate their predictive capacity to detect cognitive impairment in individuals with bipolar disorder (BD) compared to healthy controls (HC). We explore whether higher brain-PAD relates to poorer cognition in BD, as it shares neuropathological similarities with brain aging.

Specifically, we expect that all algorithms will show a stronger association of brain age with executive function in BD compared to HC. This study builds on our prior work that showed that regularized linear algorithms trained on large samples have been shown to be sensitive to BD, we expect that the algorithms such as PHOTON and the multimodal algorithm will more readily pick up cognitive differences between groups.

## Methods

### Sample

Participants were recruited from the Successful Aging Evaluation study or prior research cohorts (Dev et al., 2015) and included 44 individuals with bipolar I disorder who were in a stable mood state. Diagnosis was based on criteria from the Diagnostic and Statistical Manual of Mental Disorders, Fourth Edition, with inclusion limited to those whose first mood episode occurred between ages 13 and 30. Stability was defined by low scores on standardized clinical rating scales for depression, mania, and psychosis. Healthy comparison participants were screened using a structured diagnostic interview to rule out psychiatric conditions. All participants completed the Delis–Kaplan Executive Function System cognitive battery. Inclusion criteria required participants to be right-handed, free from neurological or substance use disorders, and eligible for magnetic resonance imaging.

Cognitive testing was conducted during the same session as scanning for a subset of participants, while blood draws were performed on a different day, with an average delay of 82 days. All procedures were approved by the Institutional Review Board of the San Diego Veterans Affairs Healthcare System.See Dev et al (2015) for complete sample details.

### Brain imaging

Imaging data were collected using a 3T GE Discovery MR750 system, capturing a variety of modalities: T1-weighted anatomical scans, DTI, CBF, task-based fMRI, and resting-state fMRI. T1-weighted images were high resolution (1×1×1 mm voxel) for cortical and subcortical analysis. DTI scanned the whole brain (2.5 mm resolution) to assess diffusion, while CBF used pseudo-continuous ASL to measure perfusion. Task fMRI focused on BOLD signal during a delayed match-to-sample working memory task, and rsMRI captured resting-state connectivity with eyes open.

Preprocessing included motion and distortion corrections for DTI and rsMRI, with DTI data registered to MNI space and FA/MD maps extracted. T1 images were processed using FreeSurfer for cortical reconstruction and ROI measurements. CBF data were aligned to anatomical space and processed with AFNI, focusing on regions like the DLPFC and ACC. fMRI data underwent slice timing, motion correction, and GLM analysis for task activation, while resting-state data were processed for seed-based connectivity analysis. The dataset includes features across cortical thickness, subcortical volumes, FA/MD, CBF, and resting-state functional connectivity. Refer to Chapter 2 for more details.

### Brain age algorithms

We conducted analyses on brain-PADs from four brain age algorithms: (1) PHOTON, a ridge□regression model trained on FreeSurfer□derived cortical thickness and subcortical volumes from 2,188 HCs (2) brainageR, a nonparametric Gaussian Process Regression model fit to PCA□reduced voxel□based T1 scans from 3,377 HCs that extracts GM/WM/CSF maps via SPM12; (3) T1 Ensemble DenseNet, which combines preprocessed volumetric T1-weighted and axial T2-weighted scans through a 121-layer convolutional network trained on ∼1500 HCs, and (4) our Multimodal algorithm, which integrates a range of pre-processed brain imaging features using ridge regression trained on 51 HCs (see Table 2.1). These were previously described in detail in Chapter 2.

### Executive function performance

Cognitive performance was assessed with the D-KEFS. Testing was carried out by trained raters in a standardized fashion according to the manual. Two main subtests of the battery are based on the Trail Making Test (Tombaugh, 2004) and the Stroop Test (Stroop 1935) From these subtests, we selected nine raw scores for further analysis: Trails Number Sequencing Completion Time (TMT-A), Trails Letter-Number Sequencing Completion Time (TMT-B), TMT B-TMT A Difference Score, Color-Word Interference Color Naming, Color-Word Interference Word Reading, Color-Word Interference Inhibition, Color-Word Interference Inhibition/Switching, Stroop Interference (Inhibition) Effect and Stroop Switching (Inhibition□+□Switch) Effect. These scores were selected because they capture core domains of executive function, i.e., processing speed, cognitive flexibility, and inhibition, that consistently emerge as distinct components in prior dimensionality reduction studies (Camilleri et al., 2021; Clark et al., 2010; Konstantinopoulou et al., 2021; Savla et al., 2012).

### Statistical Analyses

Demographics and individual executive function scores were compared between groups with ANOVA.

For each brain age algorithm, we regressed brain-PAD on age, sex, and their interaction to obtain residuals, then used those residualized brain-PADs (which we refer to subsequently as brain-PADs throughout the text) in analyses.

The nine executive function scores (See Supplemental Figure 3.1 for distribution of scores) were subjected to principal component analysis (PCA). This was done to reduce dimensionality of the executive functioning variables and is consistent with past studies that created composite measures of executive function (Crane et al., 2008; Gibbons et al., 2012). We first assessed the normality of cognitive scores using the Shapiro-Wilk test and identified and removed outliers (scores exceeding 3 SD above the mean). Subsequently, the scores were log-transformed and scaled. We conducted PCA and identified a first component. Further, to match the procedure used for the brain-PAD measures, we residualized PC1 on age, sex, and their interaction to obtain residuals, then used the residualized PC1 (which we refer to subsequently as PC1) in analyses.

To test our hypotheses, we performed a linear regression with PC1 as the dependent variable, with brain-PAD and diagnosis (BD vs HC) and their interaction term as predictors. Given the absence of any significant interactions for any algorithm, we combined the two diagnostic groups and examined the relationship of brain-PAD to PC1 for each algorithm separately. We calculated partial eta squared effect sizes using e; these were compared across algorithms.

## Results

**Table 3.1.** Executive functioning measures and demographics differences across BD and HC.

|  | Mean (SD) |  | Statistic<br>(t, otherwise<br>stated) | p |
| --- | --- | --- | --- | --- |
|  | HC<br>(N = 73) | BD<br>(N = 44) |  |  |
| Sex (Male) | 33 (45.2%) | 12<br>(27.3%) | $\chi^2(1) = 3.01$ | 0.08 |
| Age (years) | 51.8 (12.6) | 45.4 (10.25) | 2.85 | 0.05 |
| Trails Number Completion Times Raw (TMT A) | 1.45 (0.14) | 1.48<br>(0.14) | 0.77 | 0.43 |
| Trails Letter-Number Completion Times Raw (TMT B) | 1.85 (0.17) | 1.92 (0.20) | 2.03 | <0.05 |
| TMT B- A Difference Score (TMT B - TMT A) | 1.75 (0.20) | 1.80 (0.30) | 1.09 | 0.27 |
| Color Word Interference - Color Naming Raw | 1.44 (0.08) | 1.47 (0.08) | 1.94 | 0.05 |
| Stroop Interference (Inhibition) Effect | 1.42 (0.16) | 1.45 (0.17) | 0.99 | 0.31 |
| Stroop Switching (Inhibition $\square + \square$ Switch) Effect | 1.49 (0.14) | 1.54 (0.16) | 1.67 | 0.09 |
| Color Word Interference - Inhibition/Switching Raw | 1.75 (0.08) | 1.78 (0.09) | 1.52 | 0.13 |
| Color Word Interference - Inhibition Raw | 1.71 (0.09) | 1.72 (0.09) | 0.50 | 0.61 |
| Color Word Interference - Word Reading Raw | 1.33 (0.09) | 1.33 (0.09) | 0.49 | 0.61 |

**Figure.**
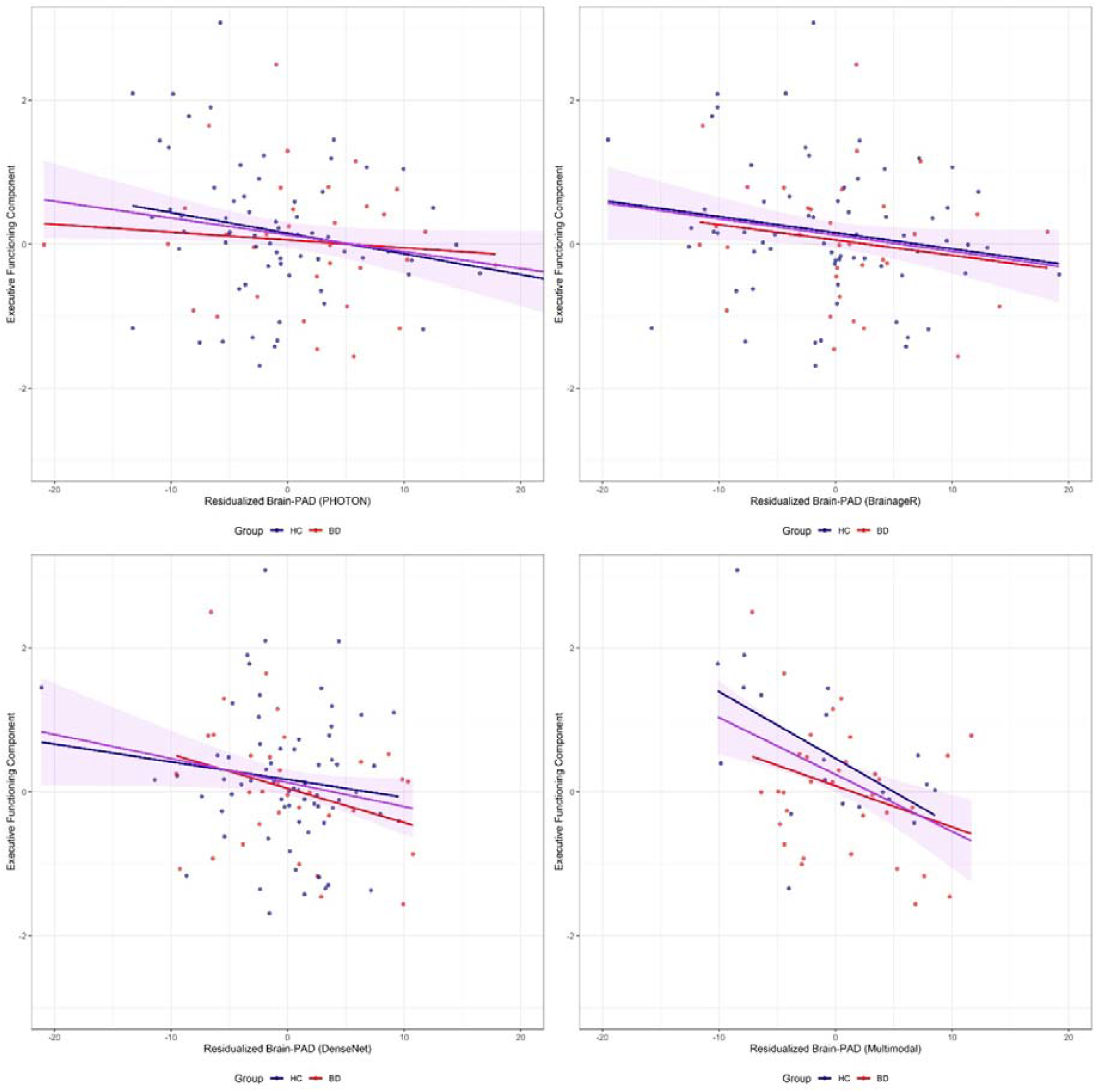
Error! No text of specified style in document..1. Relationship between brain-PAD residuals and executive function performance in BD and HC

There was no significant difference in cognitive performance (PC1) between the two groups, with mean scores of 0.05 (SD = 0.84) and 0.27 (SD = 1.04), *t*(df) = 1.39, *p* = 0.24. . Despite poorer performance on some individual executive function scores by the BD group, they did not have a significantly lower PC1 composite than HC.

Linear regression results demonstrated no interaction between group and brain-PAD, which suggests that the strength of the relationship between executive function performance and brain-PAD does not differ between BD and HC. After combining the two diagnostic groups, we found that the Multimodal algorithm’s brain-PAD showed the strongest negative effect (β = –0.084, SE = 0.024, t = –3.50, p < 0.001, R^2^ = 0.17), indicating that brains appearing older than expected scored lower on cognition. BrainageR brain-PAD also significantly predicted cognitive performance (β = –0.031, SE = 0.0116, t = –2.71, p < 0.01, partial η^2^ = 0.033, R^2^ = 0.063), while DenseNet Brain-PAD reached a modest but significant effect (β = –0.0355, SE = 0.0177, t = –2.00, p < 0.05, partial η^2^ = 0.035, R^2^ = 0.035). PHOTON Brain-PAD showed only a trend in the same direction (β = –0.024, SE = 0.0127, t = –1.92, p = 0.058, R^2^ = 0.033).

## Conclusions

Brain-PAD was a significant predictor of cognitive performance across 3 algorithms (Multimodal, BrainageR, DenseNet), while the PHOTON brain-PAD also exhibited an effect in the same direction at trend level (p = 0.058). The highest effect size was seen for the Multimodal algorithm.

Contrary to our hypothesis, the relationships between executive functioning and brain-PAD were similar in BD and HC across all four algorithms. One possible explanation is that the relationship between brain-PAD and cognitive performance may reflect general neurobiological aging processes that are shared across diagnostic groups, rather than being uniquely modulated by bipolar disorder. In other words, brain-PAD may index normative brain aging patterns, regardless of algorithm, that affect executive functioning in a similar way across both BD and HC groups. The neural mechanisms and functions that underlie cognitive decline in BD may involve disorder-specific mechanisms that are not necessarily captured by brain-PAD across algorithms, which emphasize structural age-related deviations from normative trajectories. This interpretation is further supported by findings from Chapter 2, in which we showed that the PHOTON algorithm was most sensitive to group differences in brain-PAD; yet, it exhibited the weakest and non-significant association with cognitive performance. These findings suggest that the relationship between executive functioning and brain-PAD—detected by the Multimodal, DenseNet, and BrainageR algorithms—may rely on model-specific features and patterns that are distinct from those emphasized by PHOTON. In other words, while PHOTON may be more attuned to structural deviations that differentiate diagnostic groups, these features may not be the primary factors driving individual differences in executive functioning. Conversely, the algorithms that were more sensitive to brain–behavior correlates may capture variance in brain-PAD that reflects functional or regionally distributed aging processes more tightly linked to cognition, rather than diagnosis.

Our results suggest that the multimodal algorithm may have the highest sensitivity to individual differences in executive functioning. This could be due to its inclusion of many structural and functional measures in the frontal cortex in contrast to the other algorithms that were trained on structural features throughout the brain. In other words, the Multimodal algorithm may better reflect functionally relevant brain aging processes because it integrates both functional (e.g., cerebral blood flow, resting-state connectivity, task activation) and structural information. This aligns more closely with the real-world neural demands of executive function, which rely on dynamic network coordination, not just anatomical integrity. The multimodal algorithm may be more responsive to individual-level differences that span diagnostic categories. This is especially relevant if executive function decline reflects a transdiagnostic vulnerability rather than a BD-specific deficit. The Multimodal algorithm could be more sensitive to shared aging-related or compensatory mechanisms in cognition that are shared between BD and HC. Additionally, since the Multimodal algorithm includes features heavily weighted from the frontal cortex, a region central to executive functioning, it may be better aligned with executive functioning than algorithms that distribute learning patterns across the entire brain. Finally, combining diverse range of imaging modalities and features (e.g., cortical thickness, white matter integrity, functional connectivity) might allow the Multimodal algorithm to capture interactions and latent patterns that single-modality algorithms cannot. Executive functioning is supported by distributed and interacting brain features, so the ability to detect multivariate brain aging markers may explain stronger associations between Multimodal brain-PAD and executive functioning.

However, it is also possible that the higher effect sizes for this model were due to its smaller HC training sample (N = 51) compared to the pre-trained algorithms. Algorithms trained on small sample sizes have greater epistemic uncertainty, since the combinations of imaging features across the age range is limited so the model’s predictions may be more uncertain. When the model encounters test data that appears substantially different from training data, its predictions are more uncertain as the learned patterns from the training process may not generalize, which may explain the larger brain-PAD in the BD sample in addition to true biological heterogeneity within BD and in contrast to HCs. In other words, more training data is needed to reduce the epistemic uncertainty, and disentangle aleatoric uncertainty (e.g., measurement error, natural biological variability) from it.

Prior research has also shown that regularized algorithms that are less accurate (e.g., lower MAE) may be a more meaningful biomarker for detecting pathology (More et al., 2023; Schulz, Siegel and Ritter, 2025). Generally, higher accuracy algorithms may be at risk of being overfitted at the expense of clinical utility and sensitivity to disorders (Bashyam et al 2020, Koutsouleris 2014, More et al 2023). There is growing evidence that suggests regularization pushes algorithms to focus more on features that signal atypical aging and less on features that signal typical aging as these features are down weighted; therefore, regularization increases its sensitivity to age-associated pathology, beyond patterns of healthy aging.

When examining brain-PAD’s sensitivity to cognitive performance in clinical vs. healthy samples, More et al. (2023) found that higher-accuracy algorithms better captured correlations between cognition (e.g., motor learning reaction time and Color Word Interference Test) and brain-PAD among HCs, but among individuals with Alzheimer’s, lower-accuracy algorithms showed stronger correlations between cognition (i.e., Mini-Mental State Examination scores) and brain-PAD. Additionally, More et al. (2023) also found that correlations between brain-PAD and cognitive measures disagreed when the model was applied to a test dataset from the same cohort vs. when it was applied to a test dataset from a different cohort. Cross-dataset predictions showed weaker correlations between brain-PAD and cognitive performance (e.g., TMT and CWIT inhibition time). Additionally, correcting for age bias in the within-dataset analyses showed little impact on the strength of the correlations; bias correction however, in cross-dataset correlations was less effective, only uncorrected brain-PADs showed correlations with the same cognitive scores. These results show that model application to external cohorts reduce the reliability of brain-PAD as a biomarker linking to behavior. This finding also corroborates our results, which indicate the Multimodal’s brain-PAD showed the strongest relationship (β = –0.084, t = –3.50) with the composite cognitive score.

Our study has several limitations: In this study, the Multimodal algorithm’s sample for group comparisons was relatively small (N = 22), which reduces the power to detect interactions with diagnostic group and brain-PAD from that model. Additionally, the cross-sectional design of the study precludes further investigation into consistency and reliability of brain-PAD scores and its relationship to cognitive decline over time in BD; hence we are unable to determine whether deviations in brain-PAD reflect a progressive trajectory. Longitudinal follow-up with repeated neuroimaging and cognitive assessments is necessary to characterize intra-subject changes and the clinical significance of brain-PAD over time. All participants were euthymic during the time of the scan and had BD I, so our findings may not generalize to cohorts with more severe symptoms.

Although no significant case-control differences in the relationship between brain-PAD and cognitive performance was observed in this sample, the negative relationship between brain-PAD and the composite cognitive score suggests that brain-PAD estimates may still capture clinical variation. As previously discussed, Demro et al (2022) found higher brain-PAD correlates with subclinical traits across the full sample but no differences between diagnosis groups or between first-degree relatives and controls; however, the study showed higher brain-PAD in people with psychosis compared to those without. While our study did not include samples across different psychiatric groups, our results are consistent with previous findings that subtle brain-PAD differences may emerge only when sample size (and hence power) is large enough to disentangle disorder-specific and even case-control differences. Despite these limitations, our findings contribute to the growing evidence that brain-PAD estimates, regardless of model design, can capture clinically relevant variance in cognitive function. Longitudinal and larger-sample studies are needed to elucidate the relationship of biological features and advanced aging, and clinical utility of brain-PAD in psychiatric populations.

