## Supplemental Table 1.1 and Figure 1.1 for "Cognitive Performance and Brain-Predicted Age Difference in Bipolar Disorder"

Supplementary Material

Supplemental Table 1.1 Full description of each cognitive variable

| Variable | Descriptor |
| --- | --- |
| tmtb_a | TMT B-A Difference Score  (TMT B - TMT A) |
| trnsctrw | Trails Number Sequencing Completion Times Raw (TMT A) |
| trlnctrw | Trails Letter-Number Sequencing Completion Times Raw (TMT B) |
| cwicnrw | Color-Word Interference – Color Naming Raw |
| cwiirrw | Color-Word Interference – Word Reading Raw |
| cwiwrrw | Color-Word Interference – Inhibition Raw |
| cwiisrw | Color-Word Interference – Inhibition/Switching Raw |
| stroop1 | Word Reading Raw- (Color Naming Raw +  Inhibition Raw) / 2 |
| stroop2 | Inhibition/Switching Raw - (Color Naming Raw +  Inhibition Raw) / 2 |

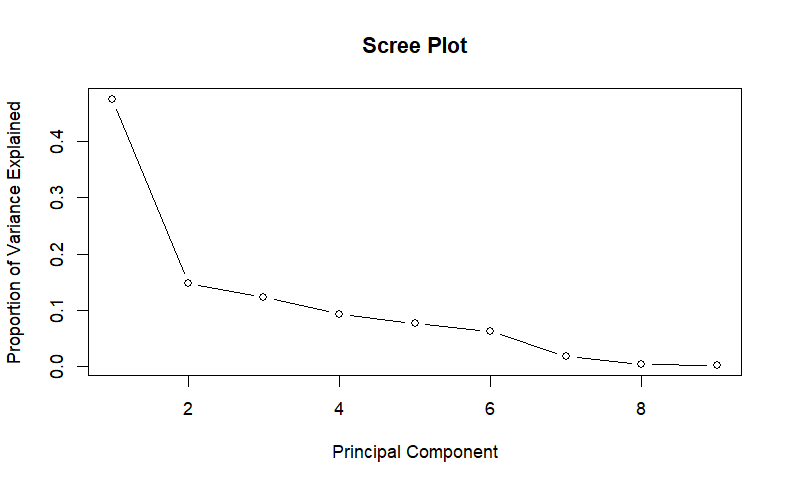

Supplemental Figure 1.1 Screeplot showing that most of the shared variance across the nine cognitive measures is captured by the first dimension (PC1).

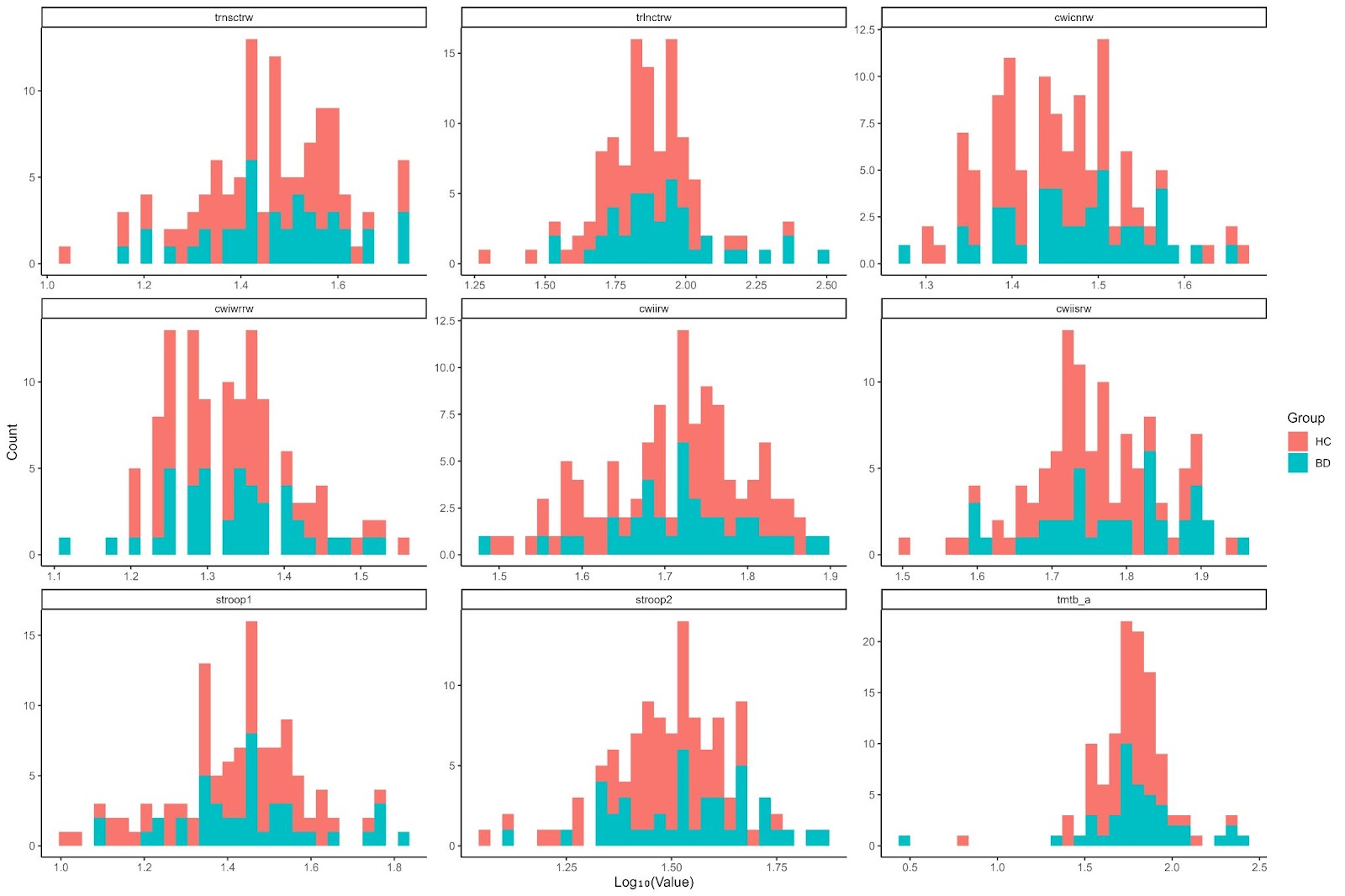

Supplemental Figure 1.2 Distribution of cognitive performance scores in HC and BD.
